# Activity Mapping the Acyl Carrier Protein - Elongating Ketosynthase Interaction in Fatty Acid Biosynthesis

**DOI:** 10.1101/2020.07.09.196451

**Authors:** Jeffrey T. Mindrebo, Laetitia E. Misson, Caitlin Johnson, Joseph P. Noel, Michael D. Burkart

## Abstract

Elongating ketosynthases (KSs) catalyze carbon-carbon bond forming reactions during the committed step for each round of chain extension in both fatty acid synthases (FASs) and polyketide synthases (PKSs). A small α-helical acyl carrier protein (ACP) shuttles fatty acyl intermediates between enzyme active sites. To accomplish this task, ACP relies on a series of dynamic interactions with multiple partner enzymes of FAS and associated FAS-dependent pathways. Recent structures of the *Escherichia coli* FAS ACP, AcpP, in covalent complexes with its two cognate elongating KSs, FabF and FabB, provide high-resolution detail of these interfaces, but a systematic analysis of specific interfacial interactions responsible for stabilizing these complexes has not yet been undertaken. Here, we use site-directed mutagenesis with both *in vitro* and *in vivo* activity analyses to quantitatively evaluate these contacting surfaces between AcpP and FabF. We delineate the FabF interface into three interacting regions and demonstrate the effects of point mutants, double mutants, and region delete variants. Results from these analyses reveal a robust and modular FabF interface capable of tolerating seemingly critical interface mutations with only the deletion of entire regions significantly compromising activity. Structure and sequence analysis of FabF orthologs from related type II FAS pathways indicate significant conservation of type II FAS KS interface residues and, overall, support its delineation into interaction regions. These findings strengthen our mechanistic understanding of molecular recognition events between ACPs and FAS enzymes and provide a blueprint for engineering ACP-dependent biosynthetic pathways.

## INTRODUCTION

Fatty acid biosynthesis (FAB) is a primary metabolic pathway in prokaryotes and eukaryotes, serving as a central biosynthetic hub for precursors utilized in membrane development and homeostasis, energy storage, cofactor biosynthesis, and signaling.^1–8^ FAB involves the iterative condensation and reduction of two carbon acetyl-CoA derived units, with each round of chain elongation requiring four separate chemical reactions. (Figure 1a).^1,3^ The first step of each cycle is carbon-carbon bond formation *via* a ketosynthase (KS) mediated Claisen-like condensation reaction between two ketide units to produce a β-keto intermediate. Subsequent reactions by the ketoreductase (KR), dehydratase (DH), and enoylreductase (ER) catalyze overall reduction of the β-position before another round of chain extension occurs. The central player in this process is the acyl carrier protein (ACP) that shuttles thioester-linked pathway intermediates to each respective enzyme active site.^9,10^ The ACP is posttranslationally modified with a prosthetic 4’-phosphopantetheine arm (PPant) at a conserved serine residue that provides the free thiol moiety to ligate pathway intermediates via a thioester bond.^11^ During ACP-mediated substrate delivery, productive protein-protein interactions (PPIs) between the ACP and partner enzymes are required.^10,12,13^ The enzymatic activities and chemical transformations in fatty acid synthases (FAS) are generally conserved, but their structural organization can be separated into two classes. Metazoans and fungi possess large megasynthases with individual domains having distinct catalytic functions (type I FAS),^14–18^ whereas bacterial and plant plastid FASs are expressed as discrete enzymes corresponding to specific genes (type II FAS).

**Figure 1.**
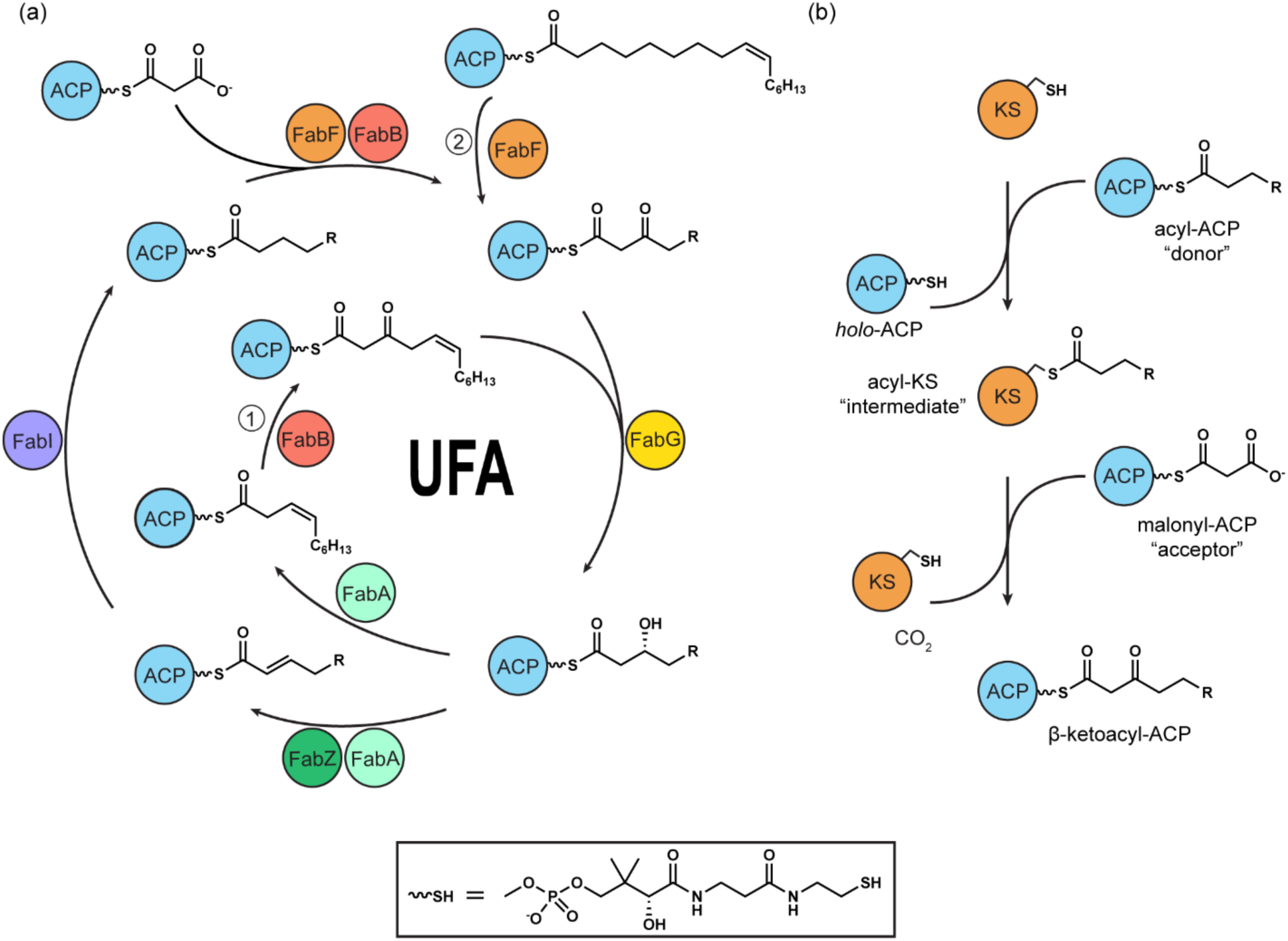
**(a)** Overview of the fatty acid biosynthetic cycle in *E. coli* focusing on the elongating KSs, FabF and FabB, and their roles in specific FAS branchpoints. For clarity, FabD-dependent production of malonyl-AcpP, cycle initiation by FabH, and PlsB/PlsC mediated chain offloading are not shown. FabA/FabB dependent branchpoint ① is responsible for the *de novo* production of unsaturated fatty acids (UFAs). Branchpoint ② is mediated by FabF and is responsible for homeoviscous adaption due to changes in temperature via the extension of *cis*-palmitoleoyl-AcpP (C16:1) to produce *cis*-vaccenic acid (C18:1). Other enzymes from FAS are: FabG, the ketoreductase (KR), FabA and FabZ the dehydratases (DH), and FabI the enoyl-reductase (ER). **(b)** Ketosynthase ping-pong reaction overview.

The most studied of the type II FAS systems is the *Escherichia coli* FAS, which has provided much of our understanding of the mechanisms and structures of FAS.^2^ In *E. coli*, biosynthesis of a single fatty acid (FA) requires that the ACP, AcpP, interacts with five protein partners during each round of chain extension. Commonly, seven rounds of chain extension occur before the acyl chain is offloaded by an acyltransferase (Figure 1a). Therefore, PPIs in type II systems between the ACP and its partner enzymes (PEs) must be fast and dynamic to ensure efficient iteration of the biosynthetic cycle.^10^ Over the past decade, a number of crystal structures of AcpP in complex with several PEs have been reported,^19–22^ and in some cases facilitated by employing phosphopantetheinamide analogs designed to ensure mechanistically-compatible crosslinking ^23–27^ to covalently trap ACP-PE complexes.^28–32^

KSs catalyze the carbon-carbon bond forming reactions that commit substrates/intermediates for each round of chain extension during FAB. Furthermore, KS substrate specificity plays a central role in the FA profile of each organism.^33^ The essential role of KSs in FAB also makes them attractive targets for the development of antimicrobials, such as platensimycin, a promising broad-spectrum antibiotic.^34–38^ *E. coli* FAS utilizes three KSs, FabB, FabF, and FabH, two elongating KSs and an initiating KS, respectively.^1,39–41^ FabH initiates *E. coli* FAB by condensing acetyl-CoA with malonyl-AcpP, while all subsequent rounds of carbon-carbon bond formation are catalyzed by the elongating KSs, FabF and FabB, using only AcpP-tethered substrates (Figure 1).^1,10^ FabF and FabB have similar substrate preferences^39,42^, but early work by Vagelos, Cronan, and Rock demonstrated that FabB is required for the *de novo* production of essential unsaturated fatty acids (UFAs).^43,44^ FabF plays an important role in homeoviscous adaptation by increasing levels of *cis*-vaccenate-containing lipids in the *E. coli* plasma membrane in response to reductions in temperature (Figure 1a).^45–47^ Despite these specialized roles in some cases, KSs share a two-step, ping-pong kinetic mechanism to condense malonyl-AcpP with acyl-AcpP (Figure 1b). In the first half-reaction, acyl-AcpP binds to the KS and delivers its thioester-bound cargo to the KS active site cysteine, producing an acyl-KS intermediate. In the second half-reaction, malonyl-AcpP binds to the KS and undergoes a decarboxylative Claisen-like condensation reaction with the acyl-KS intermediate, to produce a β-ketoacyl-AcpP (Figure 1b). During this process, the KS must undergo two separate ACP binding events and distinguish between similar, yet chemically distinct, AcpPs.

Recently, we crystallized FabF and FabB as covalent complexes with AcpPs bearing reactive fatty acyl mimetics. These structures have provided significant insight into KS-substrate recognition and AcpP-KS interactions.^29,31^ The AcpP-KS interface can be delineated into three specific interacting regions, with two flanking regions comprised predominantly of electrostatic interactions and one central hydrophobic patch.^31^ Here, we present thorough residue-by-residue and region-by-region analyses to quantify the formation of productive AcpP-KS interactions. We employed site-directed mutagenesis of the FabF interfacial residues and measured the *in vitro* and *in vivo* activities of these variants using three assays, two biochemical and one microbiological. Our results generally confirm the delineation of the AcpP-FabF interface into regions and reveal a robust interface tolerant of mutations. Additionally, we identify a catalytically important interface residue, Arg206 of FabF, that likely stabilizes the AcpP-FabF complex during chain flipping^48,49^ of acyl chain cargo from the AcpP hydrophobic core into the FabF active site. These findings further advance our quantitative understanding of the role of PPIs during bacterial FAB that may guide future metabolic engineering efforts and antibiotic development.

## MATERIAL AND METHODS

### Protein expression and purification

All recombinant proteins tested *in vitro* were expressed and purified as previously described (Figure S1a).^31^ Modifications on AcpP (holofication and acylation) were performed as previously described (Figure S1b).^50–53^ Additional details are given in the associated Supplementary Information (SI).

### Site directed mutagenesis

All FabF mutants were generated using the site-directed mutagenesis method developed by Liu and Naismith.^54^ All constructs were verified via Sanger sequencing (Genewiz). Primers used for mutagenesis in this study can be found in Table S1.

### FabF activity assay

The FabF assay mixture contained 50 mM sodium phosphate, pH 6.8, 50 mM NaCl, 0.5 mM TCEP, 250 μM malonyl-CoA, 10 μM C12-AcpP (wt) and 1 μM corresponding FabF (0 μM FabF for the control reaction) at 28 °C. 20 μL of each reaction was quenched by the addition of 10 μL of 0.3 % TFA (0.1% final TFA concentration) at 8 different time points to ensure accurate linear regression with at least 4 data points. Product formation was monitored by chromatographic separation using an Agilent 1100 series HPLC equipped with an Ascentis Express Peptide ES-18 column (15 cm* 4.6 mm, 2.7 μm) (Figure S2b), with a 1 mL/min flow rate and detection wavelength at 210 nm. A gradient method was used (A: ddH_2_O with 0.1% (v/v) TFA; B: HPLC grade CH_3_CN with 0.1% (v/v) TFA): 25% B, 0 min; 25% B, 2 min; 75% B, 12 min; 95% B, 13 min; 95% B, 15 min; 25% B, 17 min; 25% B, 20 min. *holo*-AcpP (2) concentrations were calculated from a standard curve (Figure S3b) and were used to determine initial reaction rates. C12-AcpP concentrations at 10 min were calculated using a standard curve (Figure S3a). Assays were conducted in triplicates.

### FabF catalytic parameters

Approximations of *K*_M_ and *k*_cat_ values were determined by linear regression of Hanes-Woolf plots. Additional details are provided in the Supplementary Information (Figure S4).

### *In vivo* FA profile assay

Whole cell *in vivo* Gas Chromatography Mass Spectrometry (GCMS) FA profile assays were conducted using a *fabF* knockout strain of *E. coli*, NRD23^55^ graciously provided by John Cronan (UIUC) containing a chloramphenicol resistance marker to replace nucleotides 251-434. FabF was cloned into a pBAD322-kanamycin (KAN) vector^56^ to afford an arabinose-inducible FabF cassette. CaCl_2_ competent cells of NRD23 were transformed with the control plasmid, carrying wt FabF, or respective FabF mutants. Three individual colonies from each respective FabF construct were picked and grown overnight at 37° C. Individual 50 mL flasks containing Luria-Bertani broth (LB broth, Millipore-Sigma), 0.01% arabinose, and 0.05 mg/mL KAN were inoculated with 1 mL of each overnight culture, representing three biologically independent experiments for each FabF construct. Flasks were shaken at 250 rpm at 30° C until reaching an OD_600_ between 0.6 and 0.8. Three independent extractions of each 50 mL flask were performed by pelleting 6 mL of LB media in glass vials. The supernatant was removed, and the cells were resuspended in 1 mL of 1 M HCl in methanol and incubated at 55° C for 30 minutes in a Dri-block heater. Each sample was extracted using 1 mL of hexanes and analyzed on a Hewlett Packard HP6890 series GC system connected to a 5973 MSD quadrupole MS (EI) using helium as the carrier gas and a HP-5MS column. Five μl of sample were injected while the oven was held a 120 °C for three minutes followed by a gradient from 120 °C to 230 °C at 4 °C/min followed by 5 minutes at 230 °C. All compounds were verified using NIST library searches and the results were integrated using GC/MSD ChemStation software and the relative percentages of each triplicate were averaged together (technical triplicates), to give one biologically independent reading. Three biologically independent readings were then averaged to produce the final reported values for each FabF construct.

### Cy2450B *fabB*(Ts) Δ*fabF* competent cells

The *fabF* knockout Cy2450B line was originally produced by Cronan by crossing *E. coli* NRD23^55^ with *fabB* temperature sensitive (Ts) using a Tn10 insertion with resistance for tetracycline. The resulting strain is a *fabF* knockout carrying a *fabB*(Ts) mutation. Therefore, Cy2450B contains only a single elongating KS, FabB(Ts), which is not functional at or above the non-permissive temperature of 42° C. CaCl_2_ competent cells of Cy2540B were prepared by first inoculating a 5 mL overnight culture of LB broth containing 100 μM oleic acid, 0.5 % Brij58, 0.1 mg/mL streptomycin, and 0.05 mg/mL tetracycline at 27° C. Oleic acid and low temperature were used for overnight growth in order to limit the selective pressure for the formation of revertants. 100 mL of LB broth containing 100 μM oleic acid and 0.5 % Brij58 were inoculated with 1 mL of Cy2450B from the overnight culture and grown at 27° C until an OD_600_ of 0.5 was reached. Cells were then pelleted and resuspended in a cold solution of 50 mM CaCl_2_ and 15% glycerol and incubated for 20 minutes on ice. The sample was again pelleted, supernatant was removed, and cells were resuspended in 2 mL of cold CaCl_2_ solution and 1 mL of 80% glycerol. Competent cells were aliquoted, flash frozen in liquid nitrogen and stored at -80 °C.

### Transformation of Cy2450B Cy2450B *fabB*(Ts) Δ*fabF cells*

In order to limit the number of revertants, Cy2450B cells were transformed with respective FabF constructs using a modified protocol, whereby after 42° C heat shock, cells were rescued using 500 μL LB media containing 100 μM oleic acid, 0.5 % Brij58, and incubated at 30° C for two hours. Cells were plated on LB containing 100 μM oleic acid, 0.5 % Brij58, and 0.05 mg/mL KAN and incubated at room temperature for two days.

### FabF *in vivo* complementation assay

Cy2450B competent cells were transformed (using the above protocol) with empty pBAD322-KAN vector (negative control), wt FabF (positive control), or FabF mutants. Cells were grown in either LB media, LB media with 100 μM oleic acid, or LB media with 100 μM oleic acid and 0.1% arabinose at both 37° C and 42° C (non-permissive temperature) for 24 hours. All cell lines grew at 37° C while growth at 42° C for FabF wt and FabF variants that complement FAB was only observed in the presence of arabinose and oleic acid. The OD_600_ of samples were taken the following day after 24 hours and compared to the OD_600_ of the pBAD322-KAN empty vector negative control. Cells that exhibited growth levels statistically different from the pBAD322-KAN negative control (p-value < 0.001) were classified as able to complement FAB.

## RESULTS

### AcpP-FabF interface and mutation strategy

The x-ray structure of the AcpP-FabF covalent complex (PDB ID: 6OKG) reveals a small protein-protein interface of ∼650 Å^2^ buried surface area dominated by electrostatic interactions between a positive patch on FabF and the negatively charged AcpP (Figure 2).^31^ The interface of the AcpP-FabF complex can be delineated into three regions based on the spatial organization and nature of the interactions (electrostatic or hydrophobic) (Figure 2d). We used an alanine-scanning approach, whereby we introduced one to three FabF mutations for each region in order to evaluate their individual and collective effects on FabF activity and bacterial growth. The resulting panel of FabF mutants spans all individual interacting residues, double mutants for specific regions, and triple mutants that eliminate entire regions.

**Figure 2.**
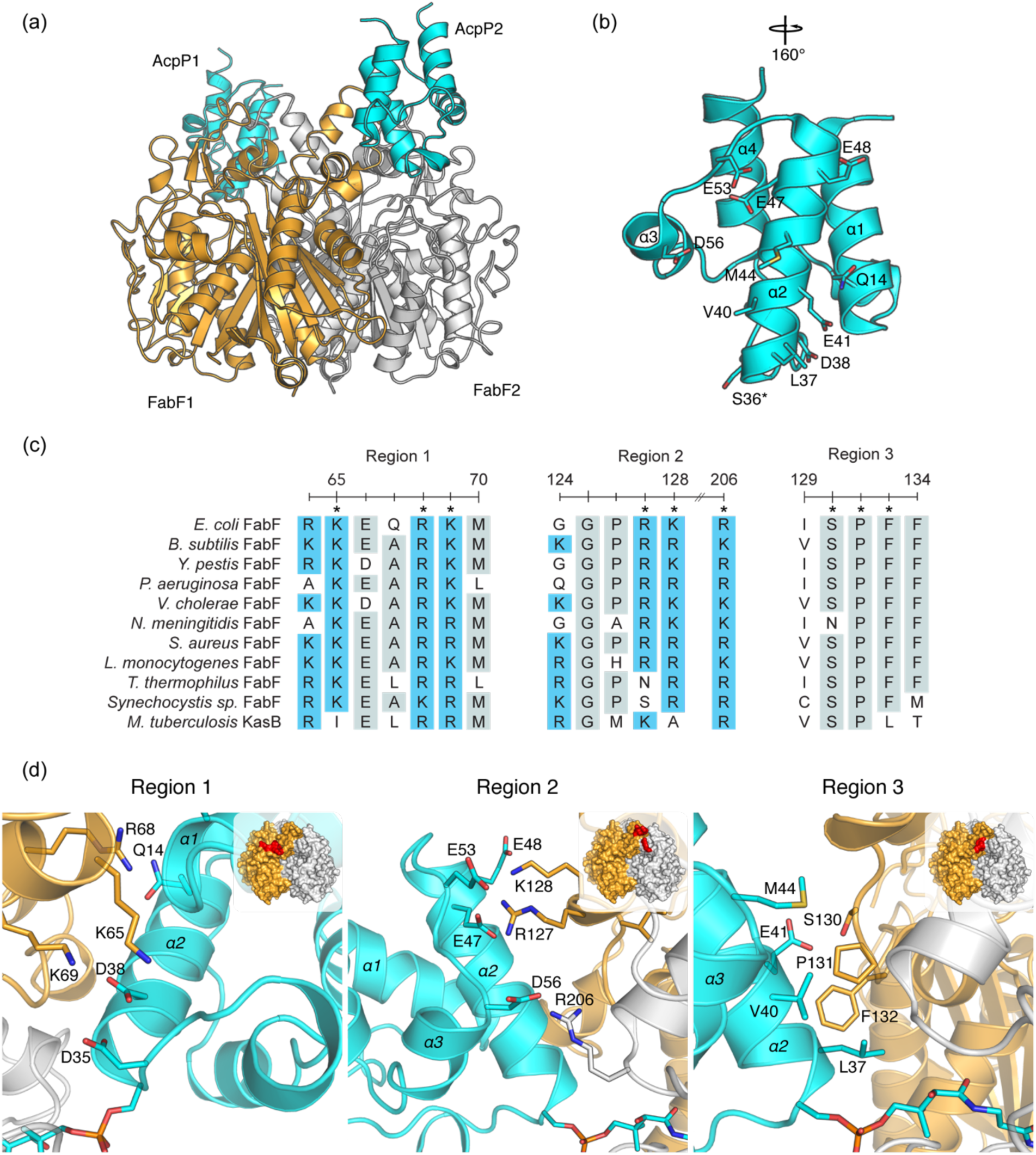
**(a)** Overview of AcpP-FabF crosslinked complex (PDB ID: 6OKG) showing a FabF dimer (orange and white) crosslinked to two AcpP monomers (cyan). **(b)** 4-helical AcpP monomer with all residues that form interface interactions with FabF shown as sticks. The conserved serine residue that serves as the attachment site for the posttranslationally installed PPant arm is marked with an asterisk (*). **(c)** Sequence alignment of *E. coli* FabF with FabF orthologs demonstrating the conservation of regions 1, 2, and 3 of the FabF interface. Positions marked with an asterisk (*) interact with AcpP in the AcpP-FabF crosslinked structure (PDB ID: 6OKG). Residues above an identity threshold of 60% are highlighted in gray, and all positively charged residues are highlighted in blue. **(d)** Interactions between AcpP and FabF shown at regions 1, 2, and 3 of the FabF interface. A cartoon inset is provided at the top right of each corner and the corresponding interface region is colored in red on the surface representation of FabF.

### FabF activity

In *E. coli* FAS, both elongating KSs, FabF and FabB, catalyze the condensation of acyl-AcpPs (donors) and malonyl-AcpPs (acceptors) to produce β-ketoacyl-AcpP intermediates (Figure 1).^42,57^ These AcpP-based reactions ensure FA biosynthesis rather than FA oxidation, the latter of which employs acyl-CoAs. However, FabF and FabB have been shown to accept CoA-based substrates, though with *K*_M_ values larger than for the corresponding AcpP-based substrates.^57,58^ Using KS substrate promiscuity for CoA-based substrates, we used an HPLC-based assay (Figure S2) using malonyl-CoA (acceptor) and C12-AcpP (donor) to interrogate FabF residues identified at the interface between AcpP and FabF (Figure 2). By monitoring the formation of *holo*-AcpP, we were able to determine the reaction rates of wt FabF and 11 FabF mutants (Figure 1, bar graph). C12-AcpP is fully converted to *holo*-AcpP by FabF wt after 10 min (Figure S2b). By analyzing the unreacted C12-AcpP at t = 10 min, we also correlated the reaction rate catalyzed by each FabF to the completion of the corresponding reaction (Figure 1, dots).

Overall, introducing single point mutations at the FabF-AcpP interface does not significantly reduce the condensation reaction rates with one exception, as the variants K65A, R68A, K128A, S130A and F132A are at least half as fast (1.37, 1.20, 2.34, 1.69 and 2.72 min^-1^, respectively) as wt FabF (2.24 min^-1^). However, R206A is six-times slower than wt (0.41 min^-1^). Both double mutants K65A/K69A (0.59 min^-1^) and R127A/K128A (1.75 min^-1^) are slower than the tested single mutants from the corresponding region. The region 1 and 2 delete variants, K65A/R68A/K69A and R127A/K128A/R206A, are both 17 times slower than wt (0.13 min^-1^), while K65A/R127A/K128A, which combines mutations from both regions, is three times faster than either region delete variant (0.40 min^-1^).The analysis of the reaction completion corroborates the kinetics results with 5 of the 6 tested single mutants (except R206A with 39% completion) and R127A/K128A fully converting C12-AcpP into *holo*-AcpP after 10 min, whereas variants K65A/K69A, K65A/R68A/K69A, R127A/K128A/R206A and K65A/R127A/K128A only partially completed the reaction (65%, 15%, 13% and 44%, respectively).

Because R206A demonstrated the largest negative effect on FabF activity of any single-point mutation, we expanded the mutational analysis of R206 to include an Asn substitution and determined the apparent catalytic parameters of the R206A/N variants and compared them to FabF wt and single variants from the electrophilic regions 1 and 2 (Table 1). While the decrease in catalytic efficiencies of K65A, R68A and K128A mainly results from an increase in their respective *K*_M app_ values, mutating Arg206 to alanine seems to have a minimal effect on the affinity for C12-AcpP, as the *K*_M app_ value is similar to that of wt FabF. Interestingly, the five-fol drop in overall catalytic efficiency of R206A is directly linked to an equivalent reduction in turnover number. This effect is observed as well when Arg206 is substituted with a polar residue. These results suggest a role for Arg206 in FabF catalysis rather than for AcpP recognition

**Table 1.** Apparent catalytic parameters of FabF wt and single variants from electrostatic regions. Reaction conditions: 28 °C, 50 mM sodium phosphate, pH 6.8, 50 mM NaCl, 0.5 mM TCEP, 250 μM malonyl-CoA, 2.5-15 μM C12-AcpP and 1 μM corresponding FabF. Apparent catalytic parameters were determined using a linear regression from Hanes-Woolf plots (Figure S4). Experiments were run in triplicate, and data shown are reported as mean ± standard error. * % efficiency is in respect to *k*_cat app_/*K*_M app_ of FabF wt.

| FabF | $k_{cat\ app}$ (min<sup>-1</sup>) | $K_{M\ app}$ (μM) | $k_{cat\ app}/K_{M\ app}$<br>(sec<sup>-1</sup>.M<sup>-1</sup>) | efficiency (%)<sup>*</sup> |
| --- | --- | --- | --- | --- |
| wt | 2.23 ± 0.08 | 1.9 ± 0.3 | 2.0. 10<sup>4</sup> | 100 |
| K65A | 1.91 ± 0.58 | 7.7 ± 0.4 | 4.1. 10<sup>3</sup> | 21 |
| R68A | 1.26 ± 0.31 | 13.6 ± 2.9 | 1.5. 10<sup>3</sup> | 8 |
| K128A | 2.15 ± 0.15 | 5.2 ± 0.6 | 6.9. 10<sup>3</sup> | 35 |
| R206A | 0.36 ± 0.04 | 1.8 ± 0.8 | 3.3. 10<sup>3</sup> | 17 |
| R206N | 0.58 ± 0.03 | 2.9 ± 0.4 | 3.4. 10<sup>3</sup> | 17 |

### Fatty acid profiles of FabF mutants

To determine how mutations to the FabF interface affect activity *in vivo*, we took advantage of FabF’s unique capacity to elongate palmitoleoyl-AcpP (C16:1) to *cis*-vaccenoyl-AcpP (C18:1) (Figure 1a).^59^ FabB performs all essential condensation reactions in *E. coli;* however, it does not produce appreciable quantities of C18:1. Therefore, we reasoned an *E. coli* cell line carrying a *fabF* knockout would present a FA profile with only trace C18:1 levels, allowing us to complement this cell line with wt *fabF* or *fabF* mutants and use C18:1 production as a biomarker for *in vivo* enzyme activity. To test this hypothesis, we utilized an *E. coli* Δ*fabF* cell line, NRD23^55^, from John Cronan (UIUC) and benchmarked its FA profile for C18:1 production via whole cell, FA methyl ester (FAME) analyses by GCMS. The FA profile of NRD23 grown at 30 °C consisted of appreciable amounts of C16:0 and C16:1, 40.9% and 51.1% respectively, but only 2.3% C18:1 (Figure 3). In contrast, previous studies report a FA profile comprised of 30% C18:1 in assays conducted at 30 °C with the *E. coli* strain, CY320, carrying wt *fabF*.^46^

**Figure 3.**
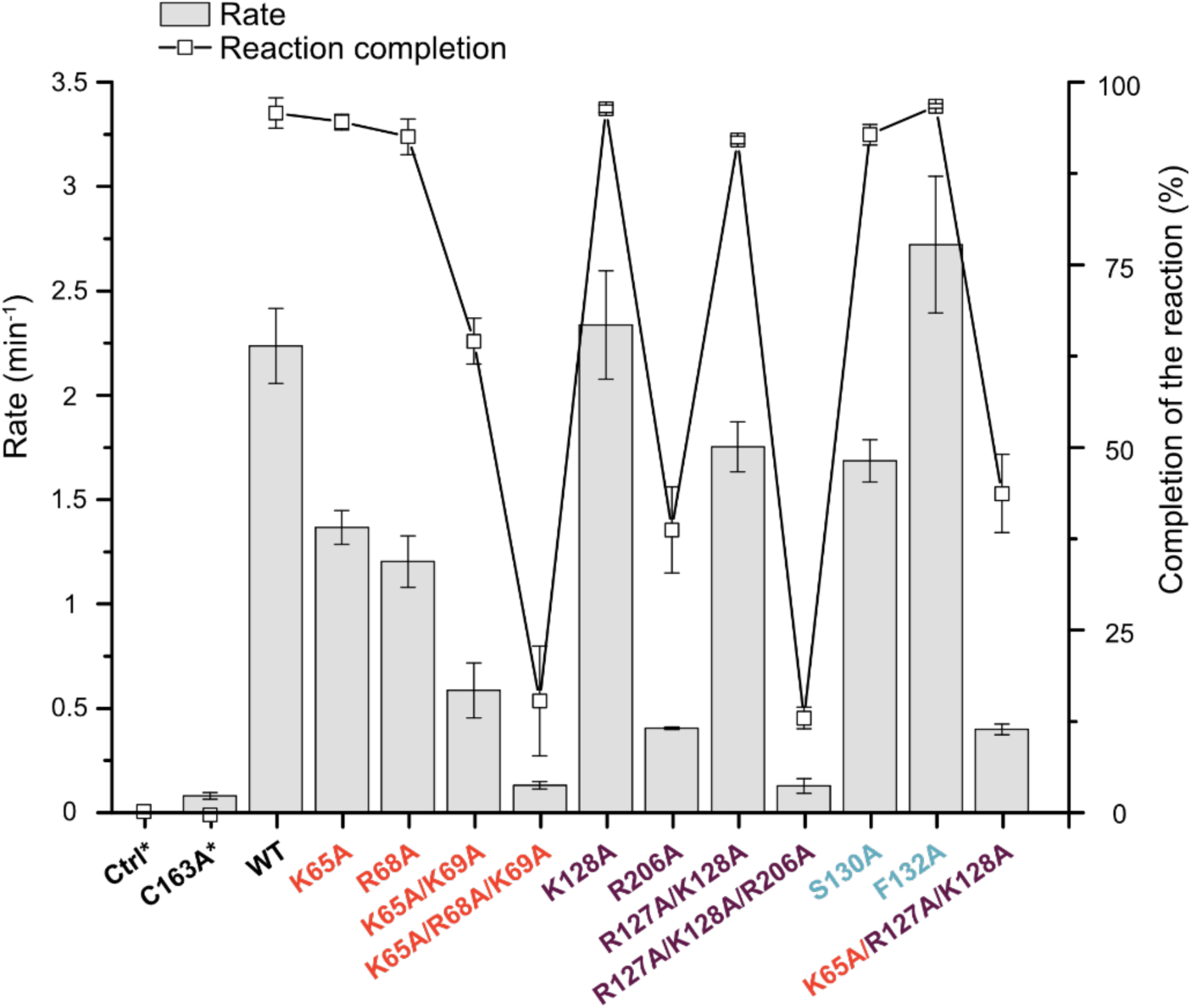
Rates of condensation reaction of C12-AcpP and malonyl-CoA catalyzed by FabF. Reaction conditions: 28 °C, 50 mM phosphate pH 6.8, 50 mM NaCl, 0.5 mM TCEP, 250 μM malonyl-CoA, 10 μM C12-AcpP and 1 μM corresponding FabF (“Ctrl” corresponds to the reaction in the absence of FabF). The completion of each reaction (0-100%) was determined after 10 min reaction time by analyzing the remaining C12-AcpP (the asterisk (*) indicates that no detectable amount of C12-AcpP was consumed after 10 min). Region 1, 2, and 3 interface mutants are indicated in dark orange, deep purple, and blue, respectively. Each reaction was run in biological triplicate and the error bars indicate the standard deviation on the mean of the three measurements.

In order to complement the Δ*fabF* strain, we cloned *wt fabF* into pBAD322, an arabinose inducible protein expression vector designed for *E. coli* FAB complementation assays.^56^ NRD23 was then transformed with either empty pBAD322 vector or pBAD322-*fabF*, and the resulting FA profiles were analyzed in the presence of arabinose to determine if an exogenously supplied *fabF* could restore C18:1 production. Indeed, our results show that induction of pBAD322-*fabF* restored the production of C18:1 and resulted in a FA profile of 35.1% C16:0, 15.8% C16:1, and 42.8% C18:1 (Figure 3). The C18:1 content in our FabF-induced NRD23 strain is higher than that of related wt background cell lines.^46^ This result is likely due to higher FabF expression levels that result from pBAD promoter-driven expression rather than the native chromosomally regulated expression of *fabF*. The resulting FA profile is more similar to the *vtr* (vaccenate temperature regulation) strain of *E. coli*, CY322, which overproduces C18:1.^46^ To further verify that the production of C18:1 was due to the induction of wt *fabF*, we generated a C163A catalytically inactive FabF mutant and assayed its FA profile. Results from the expression of FabF C163A mirrored that of the control vector and showed no production of C18:1.

We next analyzed the alterations in FA profiles of single, double, and triple mutants of the three FabF interface regions. Region 1 alanine mutants, K65A, K69A, and R68A, all showed similar changes in FA profiles, with C18:1 levels of 33.1%, 30.2%, and 35.4%, respectively, representing 22%, 29%, and 17% decreases in C18:1 content, respectively, as compared to wt FabF. To determine if the measured decreases in activities are related to charge loss at the interface, we prepared two charge-swapped mutants, K65E and K69E, and subjected them to the *in vivo* assay. Both K65E (16.4% C18:1) and K69E (15.9% C18:1) show significantly larger drops in C18:1 when compared to their alanine counterparts. In addition, the C18:1 levels in the charge-swapped mutants are similar to that of the double mutant K65A/K69A (14.6 % C18:1, which is a 69% drop from wt FabF levels). We next measured the FA profile of the region 1 delete mutant, K65A/R68A/K69A. The resulting FA profile is most similar to that of the negative control with 3.6% C18:1 content, representing a 92% drop from wt FabF levels.

Extending our analysis to FabF’s region 2, we observed similarly modest changes to the FA profile for single point mutants, with C18:1 levels of 28.7% and 34.6% for R127A and K128A, respectively. However, the FA profile of the R206A point mutant is more significantly altered than all other single mutants tested, with C18:1 content of 21.7%, representing a 50% reduction compared to wt FabF. Analysis of the R127A/K128A (20.3% C18:1) region 2 double mutant shows a generally comparable drop in activity as R206A. Given the retention of activity for all tested mutants of region 2, we analyzed the region 2 delete triple mutant, R127A/K128A/R206A. As seen for the region 1 delete, this mutant presents a similar FA profile to the pBAD322 negative control with a total C18:1 content of 3.7%.

We next examined the role of mutations of hydrophobic residues in region 3 of the FabF interface. Only point mutants for this region were tested and all mutants presented a FA profile similar to that of wt FabF. The FA profiles of S130A, P131A, and F132A contain 42.1%, 41.7%, and 39.1% C18:1, respectively. These results indicate that mutations of residues comprising FabF’s hydrophobic patch to alanine are generally tolerated.

### *In vivo* FAB complementation

Results from both the *in vitro* and *in vivo* assays suggest that FabF can tolerate some mutational disruption of the AcpP-FabF binding surface with measurable diminutions of kinetic parameters and *in vivo* fatty acid profiles. We next asked whether the residual FabF activities of some mutants sufficiently support FAB to afford survival and proliferation. Temperature sensitive mutant strains of *E. coli* developed by Cronan and coworkers have been used extensively to genetically and biochemically evaluate the function of FAS enzymes and regulators.^41,55,60,61^ John Cronan generously provided us with another *E. coli* strain (CY2450B) carrying a Δ*fabF* and temperature sensitive *fabB, fabB*(Ts). The resulting strain has only one functional elongating KS, FabB, which is incapable of performing essential FA chain extension reactions at non-permissive temperatures. CY2450 grows normally at 37 °C, but exhibits no growth at the non-permissive temperature, 42 °C. This growth arrest is due to an inadequate supply of specific FAs required for cell survival and proliferation at higher growth temperatures. Complementation of CY2450B with an exogenously-supplied, functional KS restores normal cellular growth at 42 °C if the KS produces the necessary FAs at rates that maintain viable flux through FAB.

We developed and benchmarked our *in vivo* complementation assay by transforming CY2450B with either the empty pBAD322 or our pBAD322-*fabF* construct to determine if we could restore growth at the non-permissive temperature. This assay requires oleic acid-supplemented growth media, as FabB is essential for the *de novo* biosynthesis of UFAs, and FabF cannot perform this function (Figure 1).^43,44^ Cell growth was restored at 42 °C in the presence of 0.1% arabinose and 100 μM oleic acid only when transformed with pBAD322-*fabF*, but not with the pBAD322 empty control vector or by expression of the catalytically inactive FabF C163A mutant gene (Figure 4). We then analyzed all of the interface mutants tested in our previous GCMS FA profiling studies to determine if they were capable of restoring *E. coli* growth at the non-permissive temperature. Interestingly, only the region 1 and region 2 delete mutants were incapable of maintaining cellular growth at 42 °C, while all other interface mutants were able to restore cell viability (Figure 4, Table S1). These results corroborate both the *in vitro* and *in vivo* assays and demonstrate that FabF interface mutations are relatively well tolerated and that complete removal of either region 1 or region 2 significantly compromises FabF’s critical role in maintaining cell growth and division under less than ideal environmental conditions.

**Figure 4.**
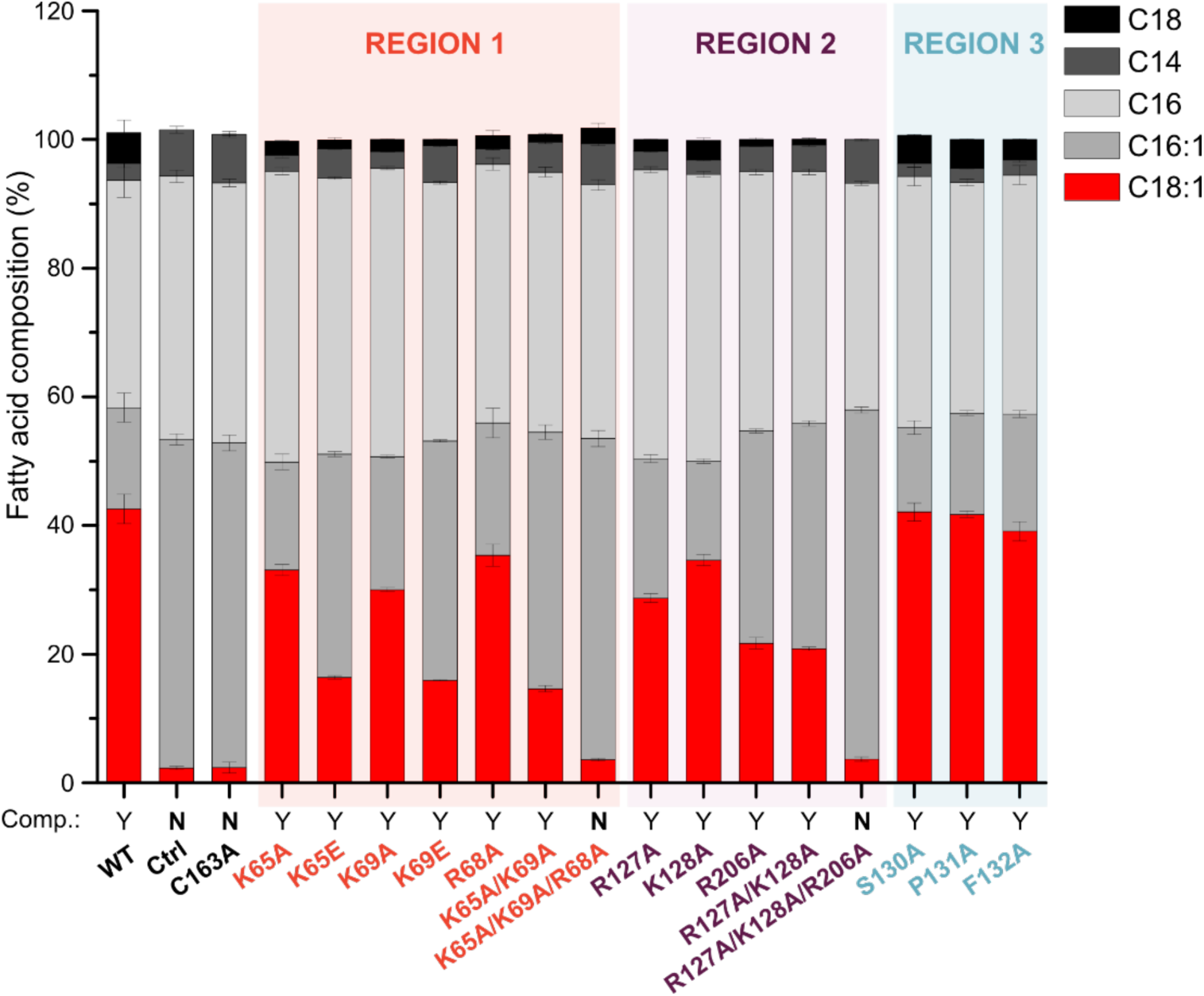
*In vivo* GCMS profile and complementation assays of FabF interface mutants. GCMS analysis of the FA profile of NRD23 Δ*fabF* strain complemented with expression of FabF interface mutants is shown in the bar chart. Total C18:1 percentage is an indicator of FabF *in vivo* activity and is shown in red. *In vivo* complementation assay results using the Cy2450B Δf*abF-fabB*(Ts) cell line are shown below the corresponding FA profile for each mutant. FabF mutants that successfully complement FAB are assigned **Y** (yes) and those that do not are assigned **N** (no). See Table S2 for raw data and statistics. GCMS data are displayed as the average percentage of total FA content. All experiments were performed as biological independent experiments in triplicate (n=3). Error bars indicate the standard deviation on the mean.

## DISCUSSION

A growing number of structures featuring AcpP in complex with PEs have recently become available, representing a significant advancement in our understanding of the complexity and diversity of AcpP-mediated PPIs.^19,20,28–31^ Moreover, these high-resolution structures, often accompanied by NMR studies and molecular dynamics (MD) simulations, provide a dynamic understanding of enzyme recognition of AcpP-tethered substrates. Despite these important advances, PPIs are complex mixtures of multiple points of interaction,^62,63^ and only a few of these studies have evaluated the importance of specific interfacial interactions involving residues on either AcpP, the PE, or both to biosynthetic pathways *in vitro* and *in vivo*.^20,29,64^

In order to catalytically map the molecular details of AcpP-KS interactions and their importance for *E. coli* FAB, we developed a multi-tier mutational strategy to assess the importance of specific residues and regions that comprise the FabF interface and subjected these mutants to both *in vitro* and *in vivo* assays. Results from these studies demonstrate that introducing single alanine mutations at the AcpP-FabF interface often have minor effects on FabF activities. These results are generally consistent with a previous analysis of the AcpP-FabB interface showing that a single alanine mutation (D38A AcpP) only modifies the *in vivo* FA profile to a modest extent.^29^ While the effects of alanine mutations are rather small, we still observe that mutations to the electrostatic regions 1 and 2 impact FabF activity more than mutations to the central hydrophobic region, emphasizing the dominant influence of electrostatics for AcpP binding. In addition, the *in vivo* activities of FabF for charge-deleted (Ala) and charge-swapped (Glu) mutants present significant differences, with the glutamate variants of region 1 producing half of the amount of C18:1 compared to their alanine counterparts. These results together indicate that the AcpP-FabF interface functions mainly through electrostatic complementarity between the acidic surface residues of AcpP and the basic residues of FabF.

However, unlike other point mutants, mutation of Arg206 to alanine resulted in a significant reduction in both *in vivo* and *in vitro* KS activity. The orientation of the guanidinium side chain of Arg206 in *apo*-FabF has been reported to inhibit CoA binding, and mutating this residue to a glycine increases FabF’s affinity for CoA-based substrates.^57^ Interestingly, when we superimpose the structures of *apo-*FabF (PDB ID: 2GFW), C12-FabF (PDB ID: 2GFY) and AcpP-FabF (PDB ID: 6OKG), Arg206 undergoes a conformational change in the AcpP-FabF structure to provide an important interaction with helix III (α3) of AcpP (Figure S5). AcpP’s α3 is a dynamic helix that exhibits structural plasticity.^9,65–67^ PPIs involving α3 have been proposed to facilitate substrate chain-flipping^49^ from AcpP’s hydrophobic core to the active site of PEs.^28,65,68–71^ The crosslinked AcpP-FabA structure (PDB ID: 4KEH), along with NMR and MD data, suggest that chain-flipping is facilitated by FabA binding to AcpP’s α3.^28^ The significant drop in activity for R206A, both *in vitro* and *in vivo*, suggests a more specific function for Arg206. To test this hypothesis, we compared the effects on *k*_cat app_ and *K*_M app_ values of R206A/N with wt FabF and alanine variants of regions 1 and 2. We observed that, unlike for Lys65, Arg68, and Lys128, mutating Arg206 to alanine or asparagine only affects turnover number (*k*_cat app_) and not *K*_M app_. This result suggests a role for Arg206 in catalysis rather than for AcpP binding, namely, to facilitate chain-flipping by acting as a wedge to pry apart helices II and III and stabilize α3 in a position suitable for substrate transfer to and from the KS active site.

We analyzed the AcpP-FabF interface using combinations of double and triple mutations of FabF, focusing on the two electrophilic contact points, regions 1 and 2. Double mutants in regions 1 and 2 decrease *in vitro* FabF activity compared to single point mutants but still enable cell growth and proliferation *in vivo*. The substitution of three mutations within a region (region delete) leads to the expression of a modified FabF that cannot support FAB to levels necessary for *E. coli* growth. The activity of both FabF region delete mutants yield *in vivo* FA profiles that are similar to the profile of the catalytically inactive FabF mutant, C163A. Moreover, the *in vitro* reaction rates of these mutants are nearly 20 times slower than wt in the conditions tested. The region delete mutants result in the net removal of three interfacial interactions, leading to a net loss of three units of positive charge from the PPI. The *in vivo* and *in vitro* data indicate that there may be a correlation between charge and activity, but the triple variant K65A/R127A/K128A, which combines mutations from regions 1 and 2, remains three times more active than both region delete mutants *in vitro*. These results suggest that charge alone cannot completely explain the reduction in activity for region delete mutants. Taken together, our results suggest that the AcpP-FabF interface evolved in a modular fashion, where only the full disruption of a region leads to an inactive FabF. This concept has been previously discussed for larger protein-protein interfaces, and our study indicates that it may be applied for the small, dynamic interface between an ACP and a PE.^62,63^

To establish whether the delineation of the AcpP-FabF interface into specific regions can be broadly applied to elongating type II FAS KSs, we aligned and analyzed sequences and crystal structures from all FabF and FabB homologues currently available in the Protein Data Bank. This analysis shows that the primary sequences of type II FAS KSs, are quite conserved for regions 1, 2, and 3 (Figure 2c, S6-7). Some of these regions contain one additional or fewer positively charged residue, or their positions vary slightly in the primary sequences, but their relative placements in interaction regions appears to hold. Alignments of region 2 show that Arg127 or Lys128 can be functionally substituted into an earlier position, namely position 124 (FabF numbering), and still maintain the spatial positioning at the interface to be defined within that region (Figures 2c,5). Similarly, Arg206, which is a conserved basic residue among FabF orthologs, is not present in FabB (Figure 5). Instead, FabB possesses Arg45, that, while in a different position in primary sequence, maintains the same relative spatial positioning at the PPI (Figures 5, S7). In the crosslinked AcpP-FabB structure, FabB’s Arg45 engages Asp56 on α3 of AcpP in a manner similar to that of Arg206 from FabF. Therefore, Arg45 likely plays an important role in AcpP mediated cargo delivery to FabB’s active site.^29^ Type II FAS KSs are known to be viable targets for antibiotic-based inhibition, as shown by the discovery and development of platensimycin.^36,37^ The conserved nature of type II FAS KS interfaces may provide an alternative strategy for targeting the KS, namely through the development of PPI inhibitors designed to disrupt the AcpP binding site.^72–74^

**Figure 5.**
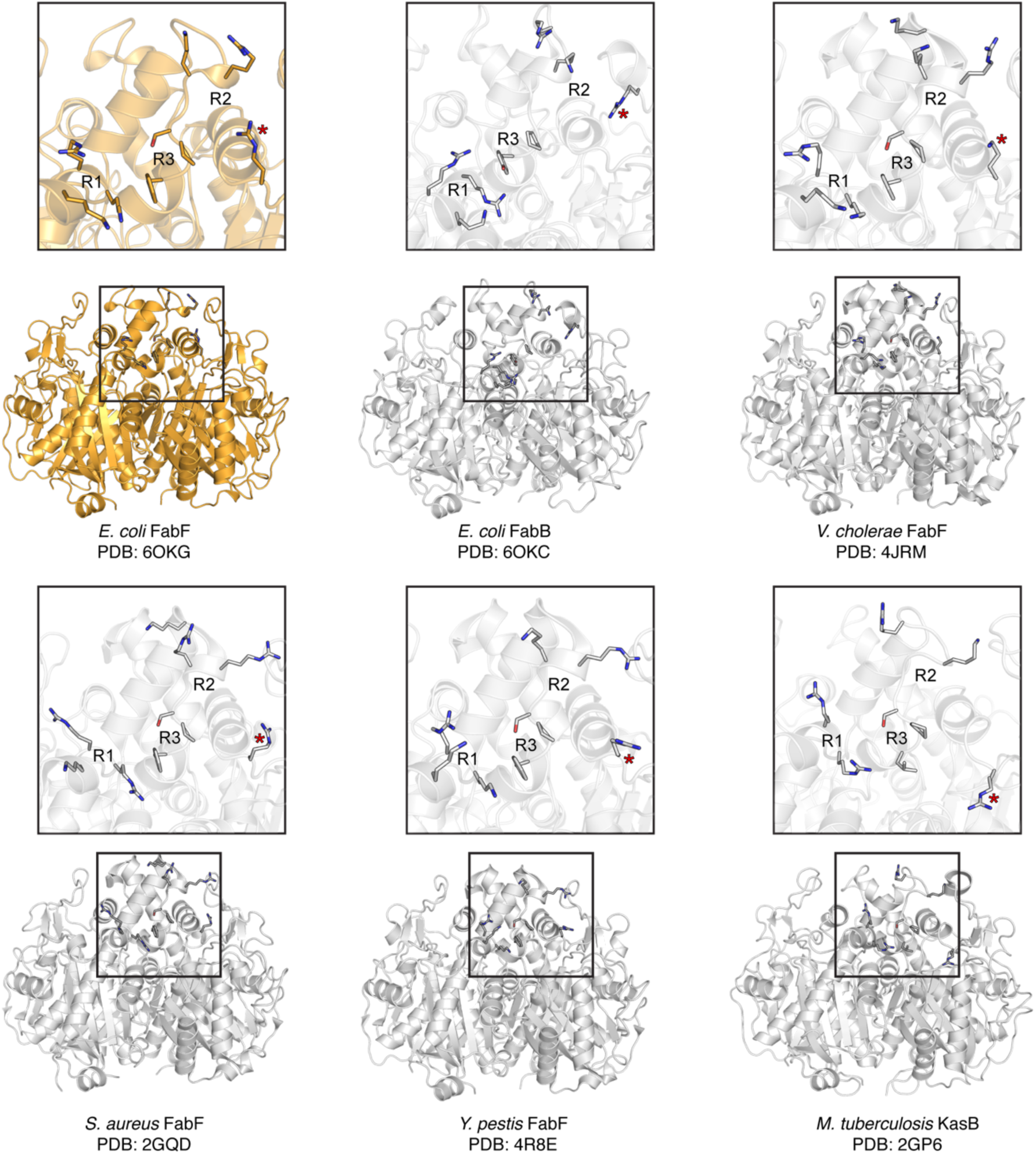
Comparison of *E. coli* FabF interface residues with that of *E. coli* FabB and four FabF orthologs from pathogenic organisms. The overall architecture of the elongating KS interfaces is generally conserved in sequence (Figure 2c) and in space with regions 1 and 2 consisting of between two and four positive residues, while region 3 is generally conserved in all FabF orthologues. A more complete structural comparison of the *E. coli* AcpP-FabF interface residues with other FabF orthologues, FabB homologues, and type II PKSs is provided in Figures S6, S7 and S10.

Given that FabF is considered the evolutionary progenitor of the KS domains found in type II PKSs,^75,76^ we aligned and compared the AcpP-FabF complex to the recently published crosslinked ishigamide ACP-KS/CLF (chain length factor) complex (Iga10-Iga11/Iga12),^77^ to determine what, if any, elements remain from the type II FAS PPI recognition motifs. A general overview of the two complexes shows that AcpP and Iga10 (igaACP) engage their respective KSs at the same binding interface, but igaACP is rotated with respect to AcpP in the AcpP-FabF structure (Figure S8). The twisted binding mode of igaACP leads to a change in the position of α2 and α3, which results in notable differences in the interfacial interactions that stabilize these two complexes. While region 1 of the ishigamide KS/CLF (igaKS/CLF) interface shares some similarities with FabF, regions 2 and 3 have diverged significantly to yield a unique set of interactions specific for igaACP (Figure S8). The igaKS/CLF region 2/3 interactions consist of a complex network of charged, polar, and hydrophobic residues that bury a salt bridge between Arg210 of igaKS/CLF and Glu51 of igaACP (Figure S9). Removal of this buried salt bridge via mutation of Arg210 to alanine results in complete loss of product formation, indicating that this interaction network is important for complex formation and specificity. Analysis of the putative ACP binding sites on the aryl polyene synthase KS/CLF^78^ and actinorhodin KS/CLF (actKS/CLF)^79^ structures also indicates an evolutionary divergence from the conserved interface interactions seen in type II FAS KSs (Figure S10). It is not surprising to find that type II FAS and PKS KSs have different interfacial interactions with their respective ACPs, as PPIs provide an elegant way to separate primary and secondary metabolic pathways in non-compartmentalized cells by discouraging pathway crosstalk.^10^ For example, the *S. coelicolor* type II FAS ACP is known to be a poor substrate for its own endogenous type II PKS actKS/CLF.^80^ The substitution of positive regions for acidic or hydrophobic regions, or for even more complex interaction networks as seen in igaKS-CLF, represents a plausible evolutionary solution to creating orthogonal pathways with internal complementarity.

There is considerable interest in manipulating PPIs as a strategy for combinatorial and metabolic engineering efforts.^81–84^ The availability of both the AcpP-FabF^31^ and AcpP-FabB^29,31^ crosslinked structures, along with the characterization of the FabF interface reported herein, provide new insights into PPI-based engineering efforts. Grininger and coworkers have successfully shown the potential of engineering KS substrate specificity and PPIs to modify the yeast FAS profile for custom biosynthesis.^85–88^ Favoring or disfavoring ACP-KS interactions by introducing mutations at the KS interface can modify chain lengths of FAs produced by the yeast FAS, although these mutants often have diminished overall enzyme activity.^88^ Similarly, Milligan et *al*. showed that expression of an AcpP D38A mutant resulted in a general reduction in the total number of UFAs in *E. coli*.^29^ While this study shows that mutating the AcpP interface can result in changes in the FA profile, AcpP interacts with multiple binding partners (Figure 1a), and any changes to the AcpP structure will likely affect associations with other PEs. The work reported herein provides an approach to manipulating type II FAS FA profiles, similar to that implemented with the yeast FAS,^88^ whereby mutation of the PE interface can selectively effect outcome of the reaction of interest. However, reductions in overall activity and flux through the FAS pathway may be a concern for implementing such a strategy. Nevertheless, mutations at the AcpP-FabF interface are well tolerated by *E. coli*, and only region delete mutations are unable to complement FAB. Therefore, KS interface mutations may serve as a viable approach to increasing the available pool of intermediate chain length acyl-ACPs for premature offloading or redirection into engineered pathways.

## CONCLUSION

In this study we applied an *in vitro* and two *in vivo* biochemical assays to systematically interrogate all interacting residues present in the recently published AcpP-FabF crosslinked crystal structure. The *in vitro* assay provided activity relationships and apparent kinetic parameters of different FabF interface mutants, while the two *in vivo* assays took advantage of KS-mediated FAS pathway branchpoints to provide a functional readout of these FabF variants. Results from these studies reveal a robust and modular FabF interface dominated by electrostatic interactions and generally support its delineation into three interacting regions. Point mutants had a minimal effect on FabF activity both *in vitro* and *in vivo*, with the exception of Arg206, which we propose to be a catalytically relevant interface residue that stabilizes AcpP’s α3 during chain-flipping events. Only the complete deletion of either electrophilic regions 1 or 2 resulted in a FabF variant that cannot sustain turnover rates capable of supporting cell survival. Sequence and structural comparison of FabF orthologs indicate that the type II FAS KS interface is generally conserved. However, the putative interfaces of KS domains found in type II PKSs have diverged, which likely serves as an evolutionary mechanism to maintain pathway orthogonality between primary and secondary metabolism in non-compartmentalized cells. The results reported herein provide a deeper, fundamental understanding of PPIs in type II FAS and provide a general roadmap for those seeking to engineer KS interface interactions.

## Acknowledgments

This work was supported by NIH GM095970. J.T.M. was supported by T32 GM832626. L.E.M. was supported by an Early Postdoc.Mobility Fellowship from the Swiss National Science Foundation. Portions of the work were also funded by the Arthur and Julie Woodrow Chair at the Salk Institute (to J.P.N.) and the Howard Hughes Medical Institute (to J.P.N.). The authors thank Prof. Dr. John E. Cronan for his assistance and for generously providing the 2 strains of *E. coli* used in this study. We further acknowledge Troy Bemis for the molecular weight determination of FabF and AcpP wt and variants.

