## Supplementary Information for "Activity Mapping the Acyl Carrier Protein - Elongating Ketosynthase Interaction in Fatty Acid Biosynthesis"

**This PDF file includes:**

Supplementary text

Figures S1 to S10

Tables S1 to S3

SI References

### Table of Contents


#### A. Biological Protocols

**A1.** Expression and purification of FabF (wt and mutants) and AcpP wt
**A2.** AcpP holofication
**A3.** *holo*-AcpP acylation
**A4**. Analysis of kinetic data
**A5.** Hanes-Woolf plots

#### B. Supplementary Figures and Tables

**Figure S1.** SDS PAGE analysis of FabF wt and variants and Urea PAGE analysis of C12-AcpP
**Figure S2** Reaction catalyzed by FabF monitored by HPLC
**Figure S3.** *holo*-AcpP and C12-AcpP calibration curves
**Figure S4.** Hanes-Woolf plots used for the determination of FabF kinetic parameters
**Figure S5.** Overlay of apo-FabF, C12-FabF and AcpP-FabF
**Figure S6.** FabF interface comparison to FabF orthologs
**Figure S7.** FabF interface comparison to FabB homologs
**Figure S8.** Comparison of AcpP and igaACP binding modes
**Figure S9.** Interface contacts between *iga*ACP and *iga*KS/CLF
**Figure S10.** FabF interface comparison to type II PKS KS/CLFs

**Table S1.** PCR primers used in this study.
**Table S2.** *In vivo* complementation assay results.
**Table S3.** Molecular weight of FabF variants and C12-AcpP

#### C. SI References

### A. Biological Protocols

**A1. Expression and purification of FabF (wt and mutants) and AcpP wt.** The genes encoding for FabF and AcpP were inserted in a pET28b vector with a cleavable N-terminal $His_6$-tag. The proteins (wt and variants) were expressed in *E. coli* BL21(DE3) cells. For each variant, a single colony was selected on the agar plate and grown overnight at 37 °C in 5 ml Luria-Bertani (LB) medium supplemented with 50 µg/ml kanamycin (KAN). Then, 1 L of LB medium (50 µg/mL KAN) was inoculated with the 5 mL-LB preculture and incubated at 37 °C until an optical cell density of 0.7 was reached. After induction with 0.5 mM IPTG, cells were grown for 3 h at 37 °C. Cells were harvested by centrifugation (500 RCF, 30 min) and the pellet was stored at – 20 °C. Cells were resuspended in lysis buffer (50 mM Tris pH 8.0; 300 mM NaCl; 10 % glycerol) and lysed by sonication. After centrifugation (17.400 RCF, 45 min), the supernatant was transferred to Ni-NTA-column and washed with 10 CV of lysis buffer and 2 x 10 mL lysis buffer containing 10 mM imidazole. The proteins were eluted with elution buffer (lysis buffer containing 250 mM imidazole). All FabF variants expressed with yields comparable to the wt (15 ± 5 mg of protein / L of LB). FabF wt and variants were dialyzed twice over 1 L of dialysis buffer at 4 °C (50 mM phosphate pH 6.8, 150 mM NaCl, 0.5 mM TCEP and 10 % glycerol), aliquoted, frozen in liquid nitrogen and stored at -80 °C (Figure S3a). AcpP wt (expressed as a mixture of *apo*- and *holo*- forms) was subjected to $His_6$-tag cleavage by thrombin and the resulting $His_6$-tag free AcpP was purified using Ni-NTA-column supplemented with Benzamidine Separophore (bioWorld). The resulting *apo-/holo*-AcpP was then converted to its *holo* form.

**A2. AcpP holofication.** *holo*-AcpP was prepared as previously described.[1] Briefly, AcpP (*apo*/*holo*) was incubated at 37 °C overnight in the presence of the phosphopantetheinyl transferase Sfp from *Bacillus subtilis*. Final reaction concentrations were the following: 50 mM phosphate pH 6.8, 50 mM NaCl, 0.5 mM TCEP, 12.5 mM $MgCl_2$, 1 mM coenzyme A, 0.05 mg/mL Sfp and 2.2 mg/mL AcpP. The completion of the reaction was determined by urea PAGE (Figure S3b).[2] The resulting *holo*-AcpP was further purified on a Superdex 75 HiLoad 16/60 size exclusion chromatography (SEC) column equilibrated with buffer (50 mM Tris pH 8, 50 mM NaCl, 5 % (v/v) glycerol, 0.5 mM TCEP). The eluted *holo*-AcpP was collected and concentrated using Amicon Ultra Centrifuge Filters (Millipore) with 3 kDa molecular weight cut off, then converted to its acyl form.

**A3. *holo*-AcpP acylation.** C12-AcpP was prepared as previously described.[3–5] Briefly, *holo*-AcpP was incubated for 3h at 37 °C in the presence of lauric acid and the acyl ACP-synthetase AasS from *Vibrio harveyi*. Final reaction concentrations were the following: 50 mM Tris pH 8, 50 mM NaCl, 0.5 mM TCEP, 12.5 mM $MgCl_2$, 2 mM lauric acid (prepared fresh in EtOH), 0.02 mg/mL AasS and 0.75 mg/mL *holo*-AcpP. The completion of the reaction was determined by urea PAGE (Figure S3b).[2] The resulting C12-AcpP was further purified on a Superdex 75 HiLoad 16/60 size exclusion chromatography (SEC) column equilibrated with buffer (50 mM sodium phosphate pH 6.8, 125 mM NaCl, 5 % (v/v) glycerol, 0.5 mM TCEP). The eluted C12-AcpP was collected, concentrated using Amicon Ultra Centrifuge Filters (Millipore) with 3 kDa molecular weight cut off, frozen in liquid nitrogen and stored at -80 °C.

**A4. Analysis of kinetic data.** Kinetic data were analyzed with Origin software. Data points from 10 s to 10 min were fitted by linear regression to give the initial velocity of substrate turnover.

**A5. Hanes-Woolf plots.** $1/k_{cat\ app}$ and $K_{M\ app}/k_{cat\ app}$ were determined by standard linear regression of [C12-AcpP]/rate upon [C12-AcpP] (equation 1) The standard errors (SE) for $K_{M\ app}$ and $k_{cat\ app}$ were calculated using the equations 2 and 3, respectively.

$$\frac{[\text{C12-AcpP}]}{\text{rate}} = \frac{1}{\nu_{max}}[\text{C12-AcpP}] + \frac{K_M}{\nu_{max}} \quad \text{where } \nu_{max} = k_{cat} \text{ as [FabF] = 1 } \mu\text{M} \qquad \text{eq. 1}$$

$$\text{SE}\left(\frac{K_M}{k_{cat}}\right) = \frac{K_M}{k_{cat}}\sqrt{\left(\frac{\text{SE}(K_M)}{K_M}\right)^2 + \left(\frac{\text{SE}(k_{cat})}{k_{cat}}\right)^2} \qquad \text{eq. 2}$$

$$\text{SE}\left(\frac{1}{\text{slope}}\right) = \text{SE}(k_{cat}) = \frac{1}{\text{slope}}\left(\frac{\text{SE (slope)}}{\text{slope}}\right) \qquad \text{eq. 3}$$

### B. Supplementary Figures and Tables

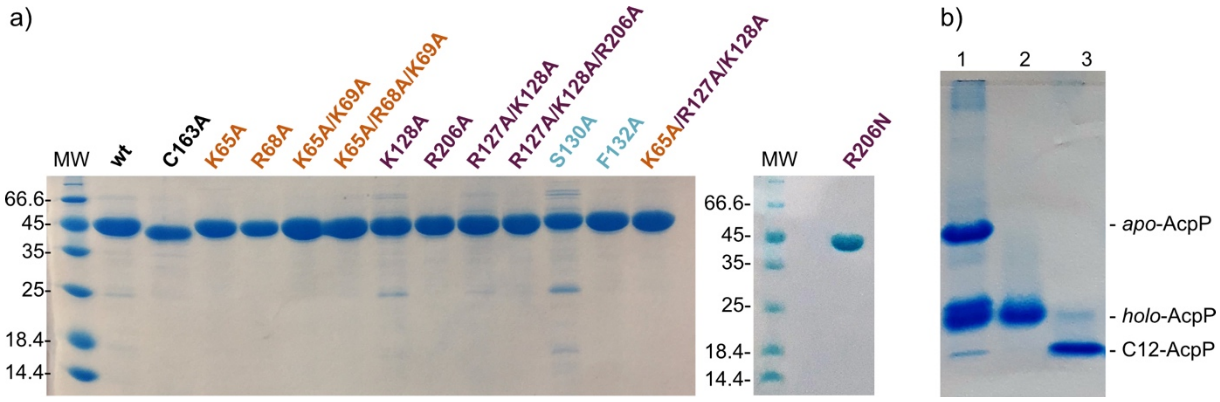


**Figure S1 a)** 15% SDS PAGE (170 V, 1 h) of FabF wt and variants, **b)** 20 % Urea PAGE (170 V, 1h30) of AcpP after purification showing a mixture of *apo*- and *holo*- forms (1), after holofication (2) and after acylation (3)

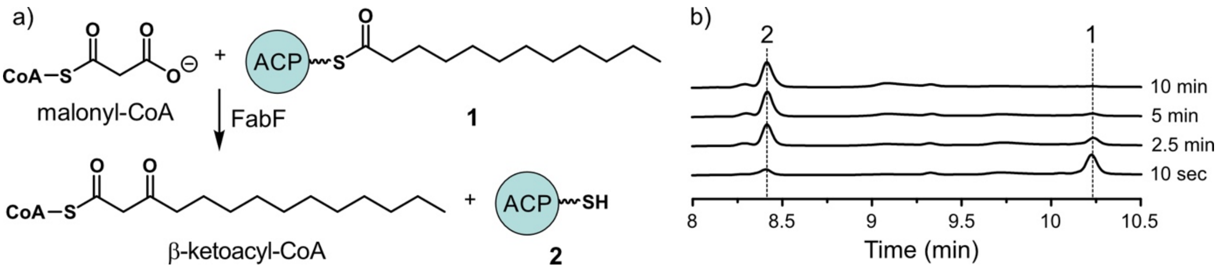


**Figure S2 a)** Condensation of malonyl-CoA and C12-AcpP (1) catalyzed by FabF. β-ketoacyl-CoA (acyl-malonyl-CoA) and *holo*-AcpP (2) are the products of the reaction, **b)** FabF condensation reaction monitored by HPLC at λ= 210 nm. Reaction conditions for this example: 28 °C, phosphate pH 6.8 50 mM, NaCl 50 mM, TCEP 0.5 mM, malonyl-CoA 250 μM, (1) 10 μM and wt FabF 1 μM.

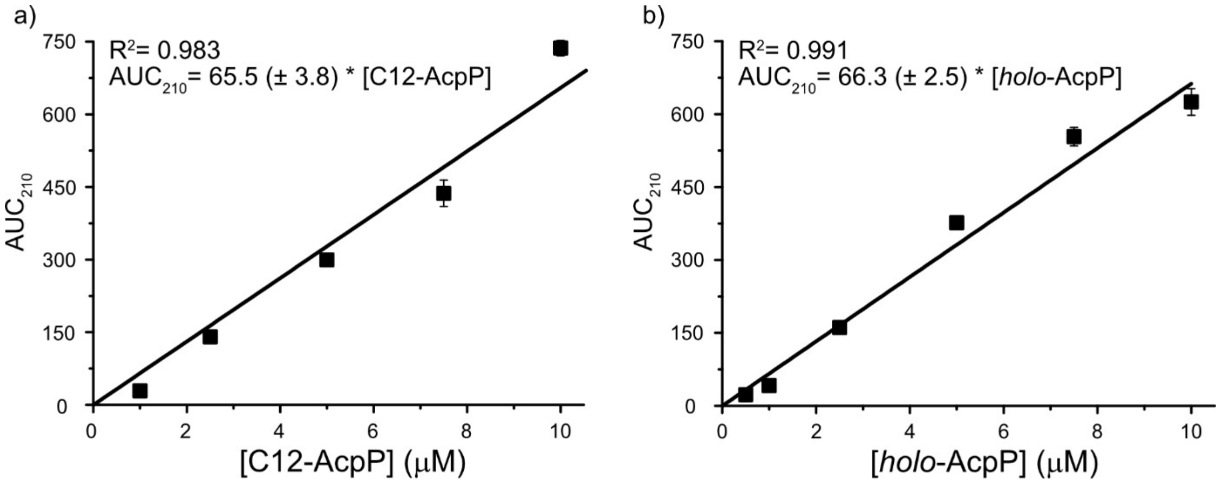


**Figure S3** HPLC calibration curves of C12-AcpP (**a**) and *holo*-AcpP (**b**). Standards (0.5-10 μM) of C12-AcpP and *holo*-AcpP were injected three times and the corresponding area under the curve (AUC) at λ=210 nm were determined. The error bars represent the standard deviation of the mean of technical triplicates.

**a**

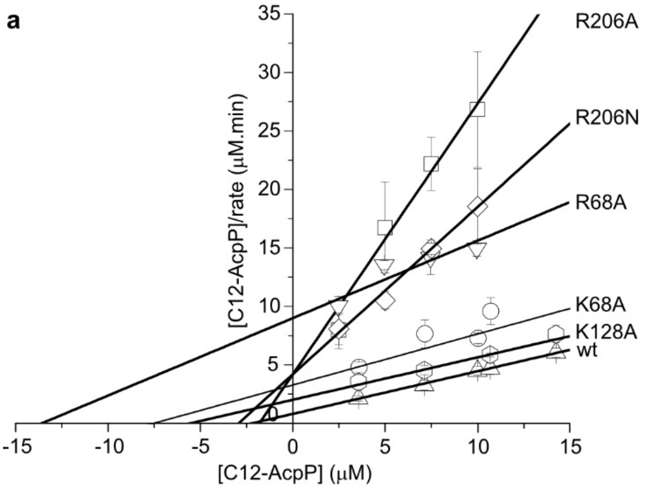


**b**

| FabF | $R^2$ | slope | intercept |
|---|---|---|---|
| wt | 0.998 | 0.365 ± 0.009 | 0.806 ± 0.073 |
| K128A | 0.970 | 0.363 ± 0.037 | 1.986 ± 0.372 |
| R206A | 0.964 | 2.314 ± 0.256 | 4.188 ± 1.942 |
| R206N | 0.991 | 1.429 ± 0.075 | 4.117 ± 0.603 |
| K65A | 0.769 | 0.436 ± 0.131 | 3.353 ± 1.021 |
| R68A | 0.837 | 0.658 ± 0.163 | 8.950 ± 1.145 |

**Figure S4 Hanes-Woolf plots of wt FabF and single variants**. **a**) Graphical representation. The error bars represent the standard deviations of the means of biological triplicates **b**) parameters from the linear regression used to determine the $k_{cat}$ and $K_M$ values of each FabF. ± indicates the standard error on the parameters.

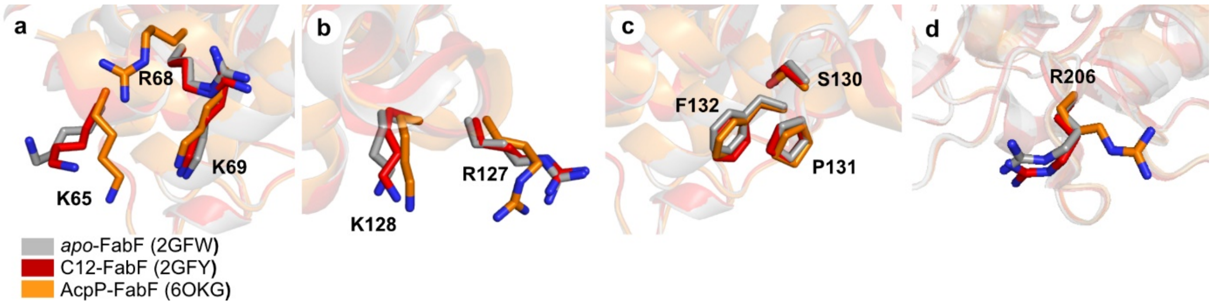


**Figure S5 Overlay of *apo*-FabF, C12-FabF and AcpP-FabF** focusing on **a)** FabF interface residues of region 1, **b)** FabF interface residues of region 2, **c)** FabF interface residues of the hydrophobic region and **d)** R206 from region 2.

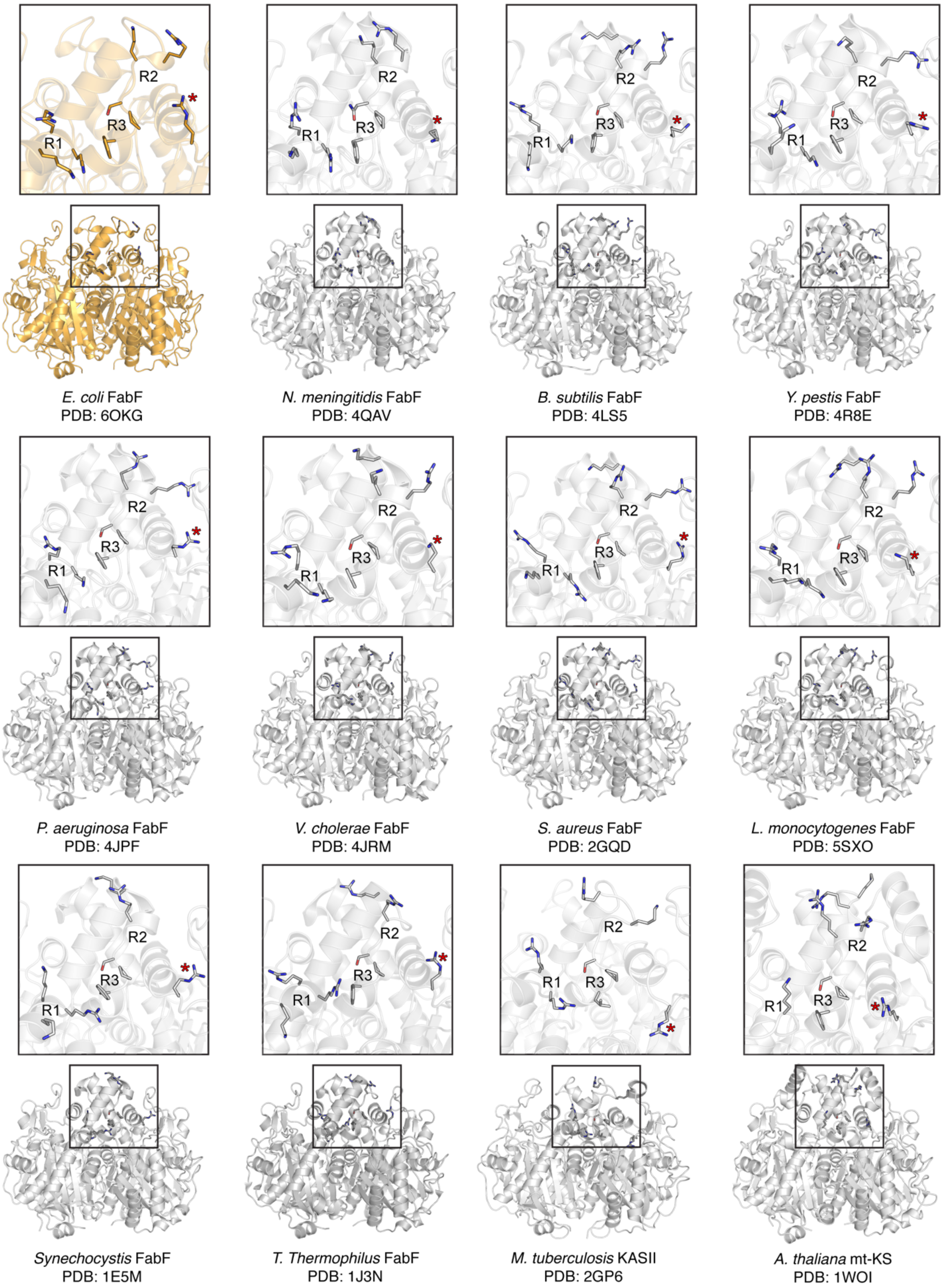


**Figure S6. FabF interface comparison to FabF orthologs.** The top left panel provides an overview of the interacting residues and designated interface regions of FabF while the following 11 panels show the putative AcpP binding residues from different FabF orthologs found in bacterial species and the mitochondrial KS from *Arabidopsis thaliana*.

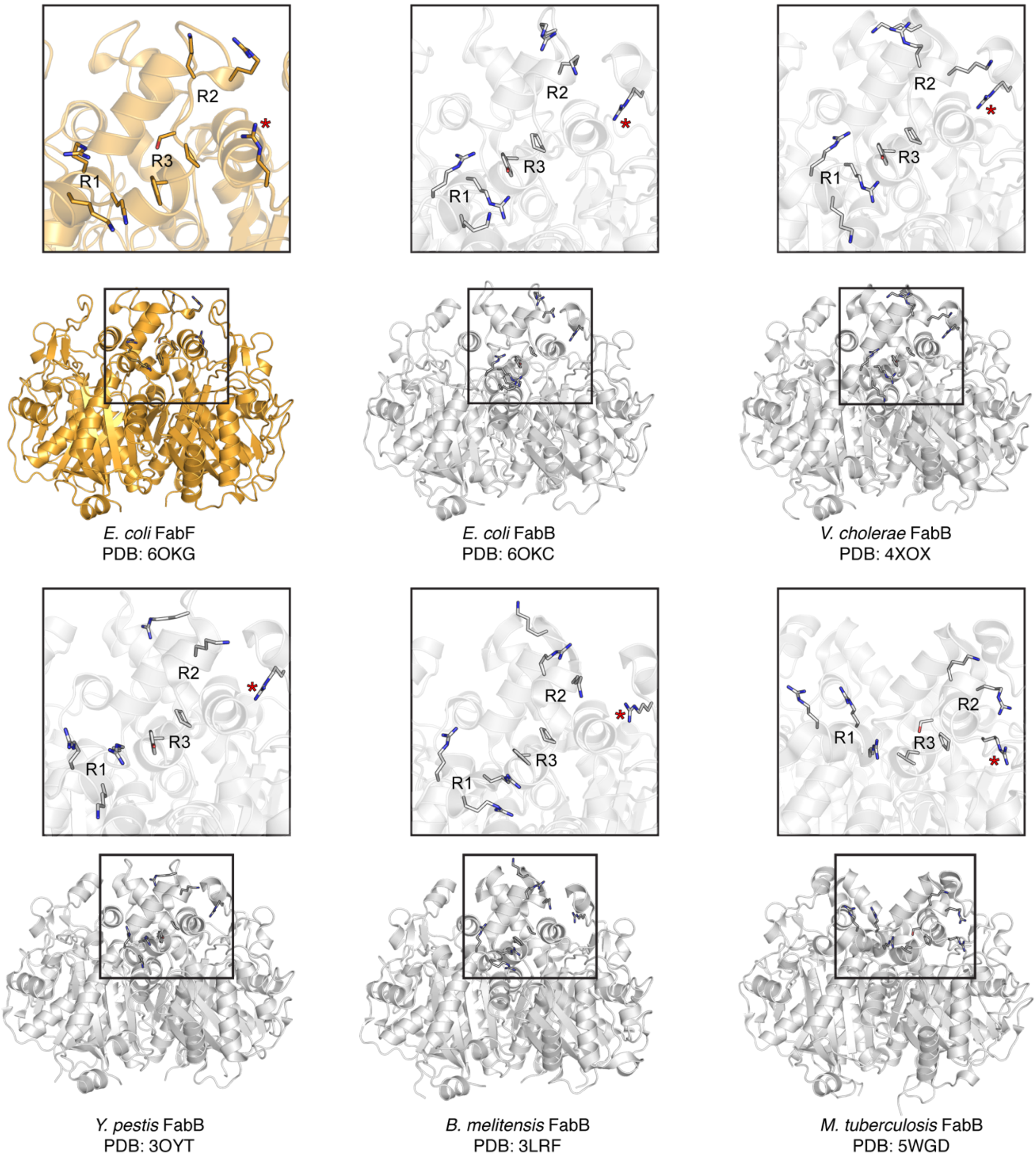


**Figure S7. FabF interface comparison to FabB homologs.** The top left panel provides an overview of the interacting residues and designated interface regions of FabF while the following five panels show the putative AcpP binding residues for five FabB homologs found in different bacterial species. The FabB Arg45 equivalent residue is marked with an asterisk (*) in each panel.

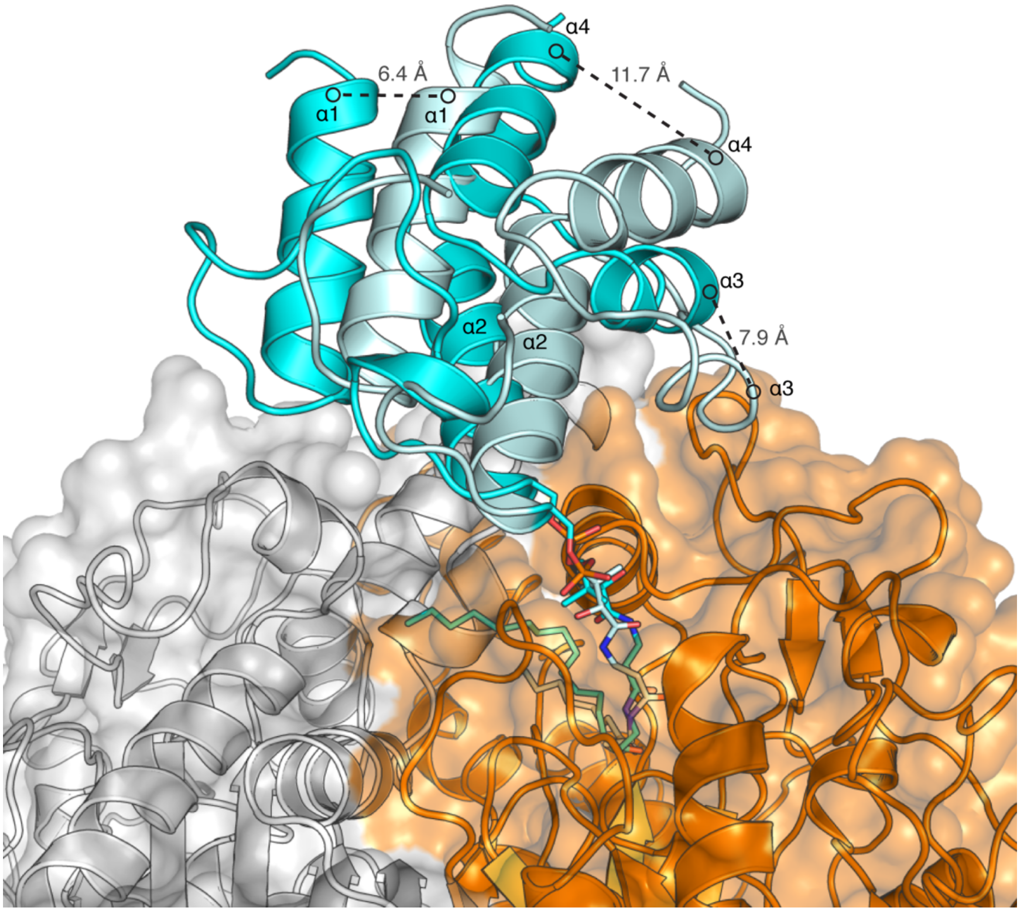


**Figure S8. Comparison of AcpP and *iga*ACP binding modes**. The KS domains of FabF and *iga*KS/CLF were overlaid in order to compare the relative binding modes of AcpP (cyan) and *iga*ACP (pale cyan). The *iga*ACP is rotated and translated with respect to the binding mode of AcpP. Measurements of the displacement are shown as dots connected by dashed lines. The KS domain to which the ACPs are crosslinked is colored orange while the other KS monomer, the CLF in the case of *iga*KS/CLF, is colored white.

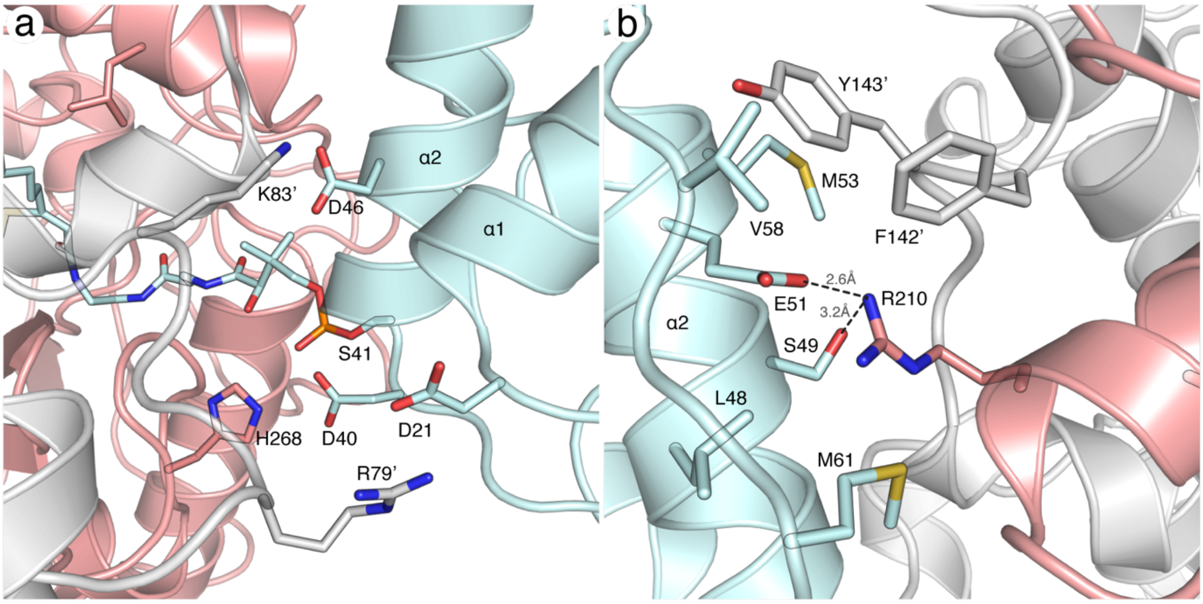


**Figure S9. Interface contacts between *iga*ACP and *iga*KS/CLF. a)** The equivalent region 1 contacts found in the igaACP-KS/CLF complex. Acidic residues (Asp21, Asp40, and Asp46) at the top of helices 1 and 2 form electrostatic interactions with positively charged residues (Arg79, Lys83, His268) on *iga*KS/CLF. **b)** Coordinated interaction network that creates a buried salt bridge between Arg210 and Glu51. The *iga*ACP is colored cyan while *iga*KS is salmon and *iga*CLF is light grey. Residues from *iga*CLF are denoted by with an apostrophe (').

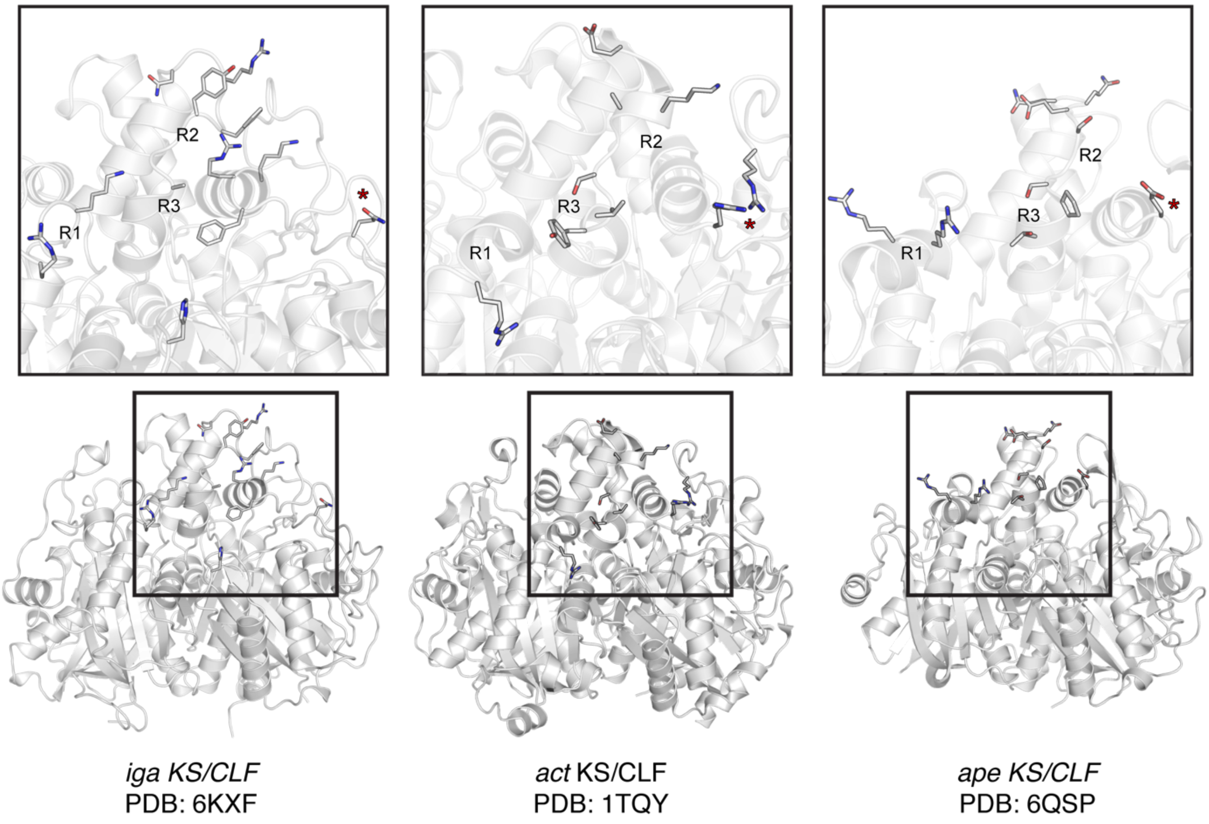


**Figure S10**. **Interface comparison of putative ACP binding sites on type II PKS KS/CLFs.** The far-left panel shows the interacting residues found at the interface of the igaACP-KS/CLF crosslinked crystal structures. The following two panels depict residues at the putative ACP binding site of the actinorhodin (*act*) KS/CLF and the arylpolyene synthase (*ape*), respectively. The interface regions are designated based on the AcpP-FabF crosslinked structure and the associated work reported herein.

**Table S1** PCR Primers used in this study. All mutants were confirmed by DNA sequencing.

| Primer name | Primer sequence 5'-3' |
|---|---|
| LM_K65A_f | CGC GCA GAA CAG CGC AAG ATG GAT GCC TTC ATT CAA TAT GGA ATT G |
| LM_K65A_r | CTT GCG CTG TTC TGC GCG CGA GAT AAT GTC CTC ACA GTT AAA ATC |
| LM_K65E_f | CGC GAA GAA CAG CGC AAG ATG GAT GCC TTC ATT CAA TAT GGA ATT G |
| LM_K65E_r | CTT GCG CTG TTC TTC GCG CGA GAT AAT GTC CTC ACA GTT AAA ATC |
| LM_R68A_f | CAG GCG AAG ATG GAT GCC TTC ATT CAA TAT GGA ATT GTC GCT G |
| LM_R68A_r | CAT CTT CGC CTG TTC TTT GCG CGA GAT AAT GTC CTC AC |
| LM_K69A_f | CGC AAA GAA CAG CGC GCG ATG GAT GCC TTC ATT CAA TAT GGA ATT G |
| LM_K69A_r | CGC GCG CTG TTC TTT GCG CGA GAT AAT GTC CTC ACA GTT AAA ATC |
| LM_K69E_f | CGC AAA GAA CAG CGC GAG ATG GAT GCC TTC ATT CAA TAT GGA ATT G |
| LM_K69E_r | CTC GCG CTG TTC TTT GCG CGA GAT AAT GTC CTC ACA GTT AAA ATC |
| LM_K65A-K69A_f | CGC GCA GAA CAG CGC GCG ATG GAT GCC TTC ATT CAA TAT GGA ATT G |
| LM_K65A-K69A_r | CGC GCG CTG TTC TGC GCG CGA GAT AAT GTC CTC ACA GTT AAA ATC |
| LM_K128A_f | ACG TGC GAT CAG CCC ATT CTT CGT TCC GTC AAC |
| LM_K128A_r | TGA TCG CAC GTG GAC CAC CGT TCA TCA GAG ATG T |
| LM_R206A_f | GCA GCG GCA TTA TCT ACC CGC AAT GAT AAC CCG C |
| LM_R206A_r | GCC GCT GCC GCG CCA AAA CCA CCA AC |
| LM_R206N_f | GCA AAC GCA TTA TCT ACC CGC AAT GAT AAC CCG C |
| LM_R206N_r | ATA ATG CGT TTG CCG CGC CAA AAC CAC CAA C |
| LM_S130A_f | ATC GCG CCA TTC TTC GTT CCG TCA ACG ATT GTG AAC |
| LM_S130A_r | TGG CGC GAT CTT ACG TGG ACC ACC GTT CAT CA |
| LM_F132A_f | CCC AGC GTT CGT TCC GTC AAC GAT TGT GAA CAT GG |
| LM_F132A_r | GAA CGC TGG GCT GAT CTT ACG TGG ACC ACC GT |
| LM_R127A_f | GGT CCA GCG AAG ATC AGC CCA TTC TTC GTT CCG TCA |
| LM_R127A_r | GAT CTT CGC TGG ACC ACC GTT CAT CAG AGA TGT GTG |
| LM_P131A_f | ATC AGC GCG TTC TTC GTT CCG TCA ACG ATT GTG AAC ATG G |
| LM_P131A_r | AAC GAA GAA CGC GCT GAT CTT ACG TGG ACC ACC GTT CAT C |
| LM_R127A-K128A_f | GGT CCA GCA GCA ATC AGC CCA TTC TTC GTT CCG TCA |
| LM_R127A-K128A_r | GAT TGC TGC TGG ACC ACC GTT CAT CAG AGA TGT GTG GT |
| LM_C163A_f | GCG ACT GCC GCG ACT TCC GGC GTG CAC AAC ATT GG |
| LM_C163A_r | AGT CGC GGC AGT CGC GAT AGA GAT GCT CGG GCC AC |

**Table S2** *In vivo* complementation assay results[a]

| | AVG OD600 | SD +/- | P-value |
|---|---|---|---|
| FabF wt | 1.017 | 0.120 | 2.733E-17 |
| pBAD322 control | 0.007 | 0.006 | NA |
| FabF C163A | 0.003 | 0.002 | 7.575E-02 |
| FabF K65A | 0.895 | 0.090 | 6.464E-13 |
| FabF K65E | 0.165 | 0.059 | 6.898E-07 |
| FabF K69A | 0.859 | 0.144 | 1.606E-10 |
| FabF K69E | 0.660 | 0.130 | 9.361E-10 |
| FabF R68A | 0.703 | 0.404 | 3.712E-05 |
| FabF K65A/K69A | 0.349 | 0.214 | 7.088E-05 |
| FabF K65A/R68A/K69A | 0.021 | 0.010 | 0.006* |
| FabF R127A | 0.420 | 0.199 | 6.959E-06 |
| FabF K128A | 0.159 | 0.037 | 3.976E-09 |
| FabF R206A | 0.38 | 0.075 | 1.145E-09 |
| FabF R127A/K128A | 0.534 | 0.029 | 1.617E-15 |
| FabF R127A/K128A/R206A | 0.016 | 0.001 | 0.016* |
| FabF S130A | 0.908 | 0.116 | 8.819E-12 |
| FabF P131A | 0.799 | 0.407 | 1.235E-05 |
| FabF F132A | 0.935 | 0.074 | 4.714E-14 |

[a]All assay results were calculated using an $OD_{600}$ reading after 24 hours of cell growth and repeated in at least biological triplicate. P-values were calculated using the Dunnet's Test comparing all results to the negative control pBAD322 empty vector strain, which represent cells unable to grow at the non-permissible temperature of 42 °C due to defects in fatty acid biosynthesis.

*P-values marked with an asterisk represent complementation results that are not statistically significant from the pBAD322 empty vector negative control using a P-value cutoff of .001.

**Table S3** Molecular weight of FabF variants and C12-AcpP. Each protein was analyzed on a LC API-SQ MS (Waters). The molecular weight of each protein ($MW_{mea}$ ± standard error) was determined using ESIprot.[6]

| **FabF** | $MW_{calc}$ (Da) | $MW_{mea}$ (Da) | **ΔMW (Da)** |
|---|---|---|---|
| WT | 45288.2 | 45290.7 (± 5.2) | 2.5 |
| K65A | 45231.1 | 45234.2 (± 5.9) | 3.1 |
| R68A | 45203.1 | 45206.4 (± 6.8) | 3.3 |
| K65A/K69A | 45174.0 | 45179.2 (± 8.4) | 5.2 |
| K65A/R68A/K69A | 45121.0 | 45125.1 (± 6.3) | 4.1 |
| K128A | 45231.1 | 45239.5 (± 5.9) | 8.4 |
| R206A | 45203.1 | 45206.0 (± 5.1) | 2.9 |
| R206N | 45246.1 | 45249.7 (± 4.7) | 3.6 |
| R127A/K128A | 45178.1 | 45177.8 (± 5.8) | 0.3 |
| R127A/K128A/R206A | 45093.0 | 45094.0 (±4.8) | 3.3 |
| S130A | 45272.2 | 45277.5 (± 5.9) | 5.9 |
| F132A | 45212.1 | 45216.3 (± 4.0) | 4.2 |
| K65A/R127A/K128A | 45121.0 | 45121.7 (± 5.5) | 0.7 |
| **AcpP** | $MW_{calc}$ (Da) | $MW_{mea}$ (Da) | **ΔMW (Da)** |
| C12-AcpP* | 9576.6 | 9575.5 (± 1.1) | 1.1 |

*The expected molecular weight of C12-AcpP ($MW_{calc}$) has been calculated considering the $His_6$-tag cleavage and the installation of the acylated PPant arm on AcpP's catalytic serine residue.
